# ANIDULAFUNGIN AND HUMAN MONOCYTES REDUCE *Candida* spp. BIOFILM WITH DIFFERENT FUNGAL BIOMASS

**DOI:** 10.1101/2025.03.03.641158

**Authors:** Paula Alejandra Icely, Sofia Carla Angiolini, Emilse Rodríguez, Cecilia Vigezzi, Cecilia Paiva, María Soledad Miró, Claudio Abiega, Juan Pablo Caeiro, Fernando Oscar Riera, Claudia Elena Sotomayor

## Abstract

**Introduction:** *Candida* spp. bloodstream infection is a frequent form of mycosis with high mortality rates. Biofilm formation is a potent virulence factor for *Candida albicans* and *Candida parapsilosis*, that confers significant resistance to antifungal agents and to the innate immune response. Echinocandins such as anidulafungin are new drugs that broaden the available therapeutic arsenal for invasive fungal infection treatment. Here we evaluated the effect of anidulafungin, human-Monocytes and combined treatment on mature biofilms of clinical isolates of *Candida albicans* and *Candida parapsilosis* with different intrinsic capacity to form biofilm, and on biofilm with low (LB) and high biomass (HB).

**Methods:** Yeasts from blood isolates were molecularly identified. Lipase and acid aspartic protease production, adhesion capacity, and ability to form biofilm were evaluated; strains were classified as Weak, Moderate or Strong biofilm forming capacity. Mature biofilm with LB and HB were incubated with anidulafungin, THP-1 cell alone or exposed to both, for 22 h. Fungal damage induced by antifungal agents and/or monocytes was determined by XTT[2,3-bis(2-methoxy-4-nitro-5-sulfophenyl) 2H-tetrazolium-5-carboxanilide] metabolic assay.

**Results:** Our results showed that anidulafungin alone could be effective to reduce mature biofilms with LB of most of the *C. parapsilosis sensu stricto* isolates, independent of intrinsic BF capacity of the strain. Anidulafungin in combination with human-Monocytes was effective in biofilm with LB of *C. albicans* and *C. parapsilosis sensu stricto* clinical isolates.

**Conclusions:** The effectiveness of ANF treatment in biofilm with LB seems to be species specific, since the antifungal agent and monocytes exhibits synergistic activity on *C. albicans* and *C. parapsilosis sensu stricto* biofilm.

## Introduction

*Candida albicans* is the most common agent of bloodstream infection in ICU patients in Latin America followed by *Candida parapsilosis*^1,2^; in the central region of Argentina, our patients with systemic infections from intensive care units (ICU) had a prevalence of 48.6% for *C. albicans*, followed by 28.6% for *C. parapsilosis*^3^ and despite the availability of potent antifungal drugs, the mortality rate of systemic *Candida* infections remains unacceptably high^2^.

Among the fungal virulence factors, the biofilm formation by *Candida* spp has been related with the progression and persistence of the infection^4^, associated with the colonization of medical devices such as intravascular catheters, prosthetic heart valves and joint replacements, or in different tissues in the host^5^. These biofilms are organized communities of microorganisms with an extracellular polymeric substance (EPS); being the most common mode of microorganisms’ growth in nature. This type of growth is more resistant to antifungal treatment and immune response than their planktonic (free-living) counterparts^6^. As effector innate immune cells, Monocytes/Macrophages (Mo/Mø) have a relevant role in the host defense against *C. albicans*^7^. They have the capacity of killing the fungus and secrete a variety of soluble factors, including cytokines and chemokines^8,9^. Although interactions between *C. albicans* and the host immune system have been well investigated for *Candida* planktonically growth, less information is currently available for such interactions in a biofilm environment. Exploring murine Mø-*C. albicans* biofilms interphase, we reported that this interaction differs depending on maturation state of biofilms (early vs. mature) and the prooxidant-antioxidant balance. Studies with Confocal Scanning Laser Microscopy (CSLM) showed planes of sessile cells with yeast and hyphal elements of *C. albicans* biofilms and Mø on the top layers of mature biofilm ^4^, confirming the ability of immune cells to penetrate in the dense structure of fungal biofilm.

Components exposed or released from biofilm matrix are involved in host immune cells activation and immune evasion^5^. Members of Sap family are highly expressed during biofilm formation, and while many are involved in blocking innate immune mechanism, others participate in the NLRP3 inflammasome activation and IL-1β production^10,11^. Also, β-1,3-glucans present in extracellular matrix or release by biofilm, can be recognized by innate receptor Dectin-1 and promote the activation of immune cells^12^.

The combination of antifungal therapy and protective function of immune cells, should play a key role in overcoming the biofilm resistant mechanism. Echinocandins such as anidulafungin (ANF) are new drugs that display favorable pharmacodynamic and pharmacokinetic characteristics and have an excellent toxicological profile^13,14^. Here we provide evidence about the effect of ANF in mature biofilms of *C. parapsilosis sensu stricto* with LB and HB, and report that the treatment with ANF in combination with human-Mo is effective in reducing fungal growth in biofilm with LB of *C. albicans* and *C. parapsilosis sensu stricto* clinical isolates.

## Material and Methods

### Microorganism

The clinical isolates were obtained from patients from ICU with Candidemia, in agreement with EORTC/MSG criteria for invasive fungal disease^15^, were identified by MALDI/TOF (Biomerieux, Argentina) and molecular methods as *C. albicans* (001 and 003 isolates) and *C. parapsilosis senso stricto* (008, 010 011 and 017 isolates)^3^. Yeast cells were grown on Sabouraud dextrose agar for 48 h of culture, centrifuged at 1000 × g, washed twice in sterile PBS, counted, and diluted to the desired concentration for the experiment^16^.

### Determination of Acid aspartic protease activity

In order to semi-quantify the acid aspartic protease (Sap) activity of the isolated strains, the agar method supplemented with BSA was used ^3,17^. SGA was added with 1% BSA (Sigma-Aldrich, USA). Suspensions of *Candida* spp. (100 μl of 1 × 10^7^ cells/ml suspension in sterile PBS) were inoculated by triplicate in wells made in agar. The plates were incubated at 37 °C in aerobic conditions for 72-96h. The enzymatic activity was observed through the formation of a halo of proteolysis around the colony and calculated as the quotient between the diameters of the halo and of the colony (*PzProt*)^3,17^.

### Determination of lipolytic activity

Rhodamine-B plaque assay was used to identify and semi-quantify lipolytic activity of strains as previously described^3,17,18^. For the activity induction, 1×10^8^ *Candida* cells were suspended in 2 ml of 0.7g% Sabouraud broth (Britania, Argentina), 0.2 ml 10% CFS and 0.05 ml 2.5% Tween 80 (Biopack, Argentina) and incubated for 72h at 37 °C. Plates with Rhodamine B agar medium (1.5% agar Britania, Argentina; 1% olive oil; 0.35% Sabouraud broth; Rhodamine B 0.001%, LTD, England) were inoculated with 150 μl of each isolate’s suspension in triplicate. After 72-96h of incubation at 37 °C in aerobic conditions, the orange fluorescence diameters of the lipase diffusion halos, were visualized by exposure to UV irradiation using a transilluminator (BioDoc-IT Systems, UV). The well and the fluorescence diameters were measured to calculate the lipase (Lip) activity (*PzLip* = fluorescence diameter / well diameter). As a negative control, 150μl of Sabouraud broth was used. The *C. albicans* strains collection ATCC 36801 (low virulence) and SC5314 (high virulence) were included as virulence factor production reference.

### Cell Surface Hydrophobicity Assay

Cell Surface Hydrophobicity Assay(CSH) was determined using the MATH (microbial adhesion test to hydrocarbons)^3^. Suspensions of the isolates (3.0 ml in PBS, at OD520 nm=0.400, UV-Vis spectrophotometer Metrolab 1200, Argentina) with added to 1.0 mL of xylene (Cicarelli, Argentina). The mixture was stirred for 1min and allowed to stand 10 min for phase separation at 37 °C, the lower aqueous phase of the sample was carefully removed using a pipette, transferred to a clean test tube and the OD520 nm of the aqueous phase was read again. The relative CSH was expressed as the percentage reduction in optical density of the aqueous phase read at 520 nm, before and after mixing with xylene.

### Biofilm formation

Different inoculums of each strain were used to obtained two type of mature biofilm classified as high (HB) and low (LB) biomass in agreement with initial use of 3×10^6^ or 1×10^5^ blastoconidia respectively. Each concentration was placed per well in 96 well plates at 37°C in a humidified 5% CO2 incubator for 48 h. Quantitative measurement of biofilm formation (BF) was assessed by XTT reduction assay ^19^ performed in triplicate for all strains and the averages and standard deviations were calculated. Strains were classified as Weak (Absorbance (A) <0.310) (WBF), Moderate (0.310<A<0.570) (MBF) and Strong (0.570<A) (SBF) biofilm producer^3^.

### Cells

The THP-1 monocytic cell line (ATCC TIB202; American Type Culture Collection [ATCC®], Manassas, VA) was used as human phagocyte source and was grown in RPMI-1640 medium supplemented with 10% SBF (NATOCOR, Argentina) heat inactivated (56ºC, 30min), 2mM L-glutamine, 50Uml penicillin, 50µg/ml streptomycin and 0.05 mM 2-mercaptoethanol in a humidified CO2 incubator at 37°C. Cell viability was determined by using the trypan blue exclusion test, was ≥96%^7^.

### Antifungal agent

The echinocandin used in this study anidulafungin (ANF) was obtained from Pfizer Pharmaceutical Group (New York, NY, USA); after dissolving this antifungal agent with culture medium, the concentration used in the experiments was 0.12mg/L because the minimum inhibitory concentration (MIC) measured by Vitek2 was 0.5mg/L.

### Experimental design and assessment of fungal damage

Planktonic cells of all the isolates were used to evaluate the effect of anidulafungin (ANF) (0.12mg/L); 1×10^5^ blastoconidia were cultured with the antifungal agent per well in 96 well plates at 37°C in a humidified 5% CO2 incubator for 48 h. The fungal viability was measured by MTT assay ^20^. Mature biofilms were then incubated in the presence of ANF (Pfizer; 0.12mg/L) alone, THP-1 cells alone or in combination of both, at effector cell: target (E: T) ratio of 1:1 at 37°C in a humidified 5% CO2 incubator for 22h ^21^. We used untreated biofilms as control of 100% growth and three replicate biofilms for each condition. After incubation, we hypotonically lysed monocytes. Biofilm condition and fungal damage induced by monocytes and/or ANF was assessed by XTT reduction assay. The results are expressed as percentage of fungal damage ^21,22^. At least three individual experiments of each conditions were carried out.

### Statisticals analyses

Experiments were carried out in triplicate. Continuous variables are expressed as the means ± standard deviation of the mean (SD) and analyzed by ANOVA followed by Bonferroni test for multiple comparisons. Student’s t test was used for comparison between two variables. Differences between the experimental groups were studied using the software GraphPad Prism 7.0 (GraphPad, San Diego, CA). A P value < 0.05 was considered statistically significant.

## Results

### Virulence factors profile of clinical isolates of Candida *spp*

First, we identified the *Candida* species and evaluated the virulence profile of clinical isolates used in the experiments. All the strains were obtained from blood culture of ICU adult patients with candidaemia, identified by MALDI TOF and characterized by molecular methods as *C. albicans* (001 and 003 isolates) and *C. parapsilosis sensu stricto* (008, 010, 011 and 017 isolates)^3^. Table 1 summarized the values obtained for hydrolytic enzymes Sap and Lip, the CSH and biofilm formation capacity of the different clinical strains. In order to compare the virulence of *Candida* spp. clinical isolates, we also included in the analysis two *C. albicans* strains collection ATCC 36801 and SC5314 as low and high virulent strain respectively^4,18^.

**Table 1:** Virulence factors profile of clinical isolates of *C. albicans* and *C. parapsilosis*.

| Isolates | Strain | Virulence factors |  |  |  |  |
| --- | --- | --- | --- | --- | --- | --- |
|  |  | Sap (Pz) | Lip (Pz) | CSH (%) | Biofilm (OD) |  |
|  |  | Mean±SEM | Mean±SEM | Mean±SEM | Mean±SEM<br>Category |  |
| SC5314 | *C. albicans* | 1.83 ± 0.27 | 2.66 ± 0.22* | 24.37 ± 11.02 | 0.890 ± 0.056 | **SBF** |
| ATCC36810 | *C. albicans* | 1.16 ± 0.20 | 1.41 ± 0.26 | 51.00 ± 12.00 | 0.227 ± 0.073 | WBF |
| C001 | *C. albicans* | 1.38 ± 0.18 | 1.79 ± 0.28 | 7.36 ± 5.87 | 0.687 ± 0.270 | **SBF** |
| C003 | *C. albicans* | 1.38 ± 0.21 | 1.66 ± 0.22 | 1.00 ± 4.58 | 0.524 ± 0.301 | MBF |
| C008 | *C. parapsilosis* | 1.33 ± 0.20 | 1.38 ± 0.16 | 6.34 ± 4.80 | 0.504 ± 0.182 | MBF |
| C010 | *C. parapsilosis* | 3.74 ± 0.27* | 1.15 ± 0.12 | 15.58 ± 6.02 | 1.054 ± 0.476 | **SBF** |
| C011 | *C. parapsilosis* | 1.30 ± 0.17 | 1.00 ± 0.23 | 39.65 ± 10.20 | 0.755 ± 0.409 | **SBF** |
| C017 | *C. parapsilosis* | 1.24 ± 0.16 | 1.63 ± 0.24 | 24.34 ± 10.85 | 0.698 ± 0.276 | **SBF** |

Sap: acid aspartic protease; lip: lipase; CSH: cell surface hydrophobicity. Biofilm category; SBF: strong biofilm formation (OD > 0.570), MBF: Moderate biofilm formation (OD between 0.310 and 0.570) and WBF: weak biofilm formation capacity: (OD < 0.310) *C. albicans* strain collection SC5314 (high virulence) and ATCC 36801 (low virulence). \*p<0.01

All the clinical isolates exhibited Sap and Lip activity; only *C. parapsilosis* C011 showed higher levels of Sap (p<0.01), and Lip levels were similar in all the strains evaluated (Table 1), although *C. albicans* SC5314 exhibited an increased lipolytic activity (p<0.01). The CSH was variable between *C. albicans* and *C. parapsilosis*. When the biofilm formation (BF) capacity was analyzed, the isolates were classified as strong (SBF)(OD>0.570), moderate (MBF) (0.310 < OD < 0.570) and weak (WFC) (OD< 0.310) producers. *C. albicans* C001 and *C. parapsilosis* C010, C011 and C017 exhibited SBF capacity, while *C. albicans* C003 and *C. parapsilosis* C008 showed MBF capacity and only low virulent *C. albicans* ATCC36810 present WBF. In both *Candida* species we identified clinical isolates with SBF and MBF capacity, independent of biofilm intrinsic structure characteristics for each species.

### Effects of Anidulafungin on Candida spp in planktonic and biofilm growth

We evaluated the effect of ANF over different forms of fungal growth such as planktonic cells or mature biofilm, by exposing them to subinhibitory concentrations of ANF 0.12 mg/ml ^23^. Planktonic yeast suspension or biofilm of both *Candida* spp, was cultured with ANF and after 22 h the fungal cell viability was evaluated by determining the mitochondrial activity present in living cells (MTT and XTT assay respectively). Antifungal activity of ANF was expressed as percentage of fungal damage and referred to untreated cell viability (absence of damage) (Figure 1).

**Figure 1.**
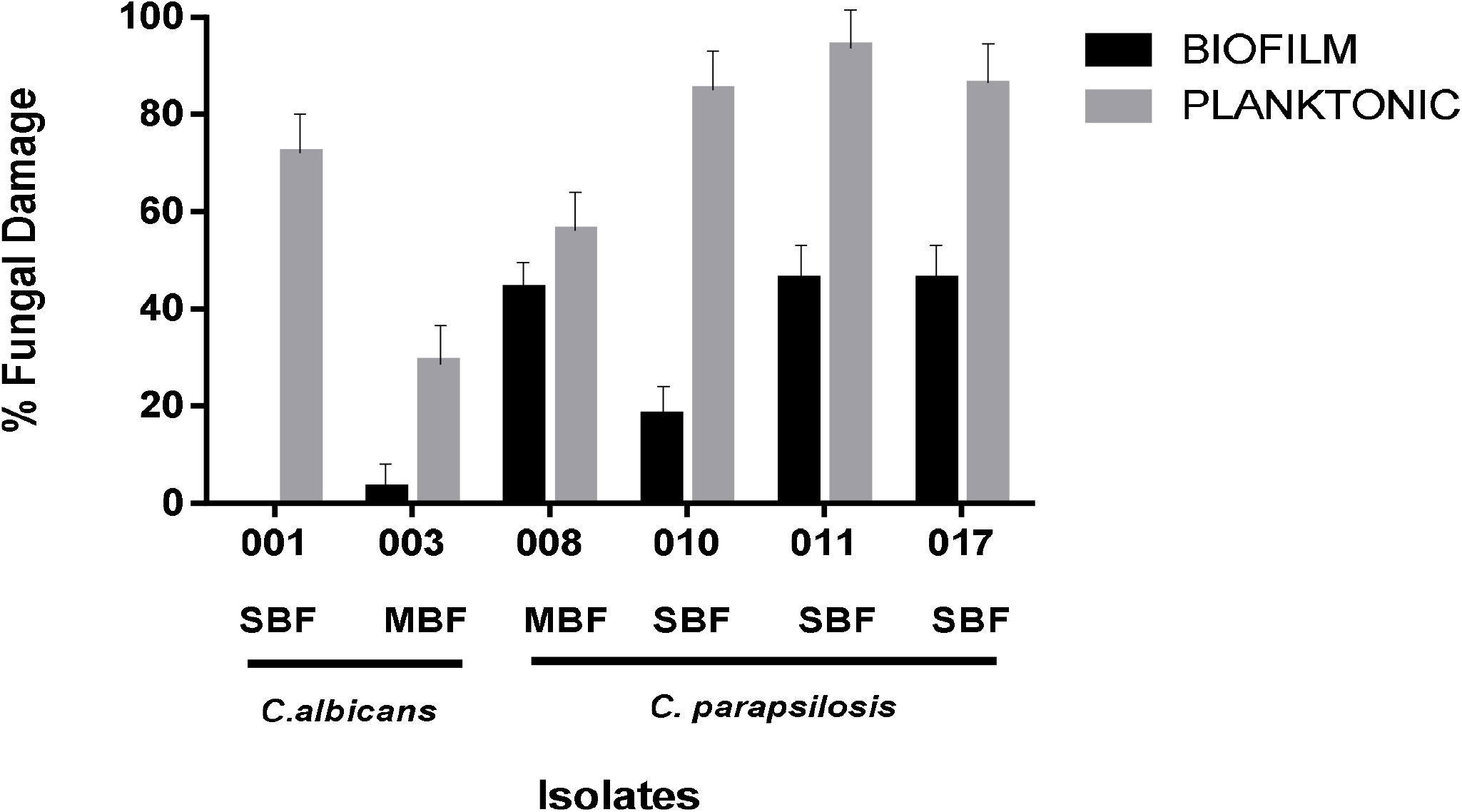
Effect of anidulafungin in the fungal growth: Percentage of fungal damage induced by subinhibiroty dose of ANF after incubation at 37°C for 22 h against *C. albicans* and *C. parapsilosis* biofilms (black bars) or planktonic cells (grey bars). The values are means± SD of three independent experiments. Each experiment was conducted using triplicate wells for each condition. The results of the damage induced by ANF was measured by XTT (Abs:495nm) or MTT (Abs: 540nm) respectively and expressed as percentage of fungal damage referred to untreated fungal cell viability (* p<0.01; **p<0.001).

On *Candida* spp. planktonic cells, ANF at the used concentration, has a strong effect against all clinical isolates tested; for *C. parapsilosis* strains the percentage of fungal damage showed values between 56-94 % (*p<0.01, **p<0.001), meanwhile in *C. albicans* isolates this parameter was 72 and 29 % depending on clinical isolate (C001 and C003 respectively; **p<0.001). As was expected, the effect of the antifungal drug wasn’t the same when the fungus grew as biofilm; interestingly for *C. parapsilosis* isolates with SBF (isolates C011, C017) and MBF (C008) capacity the ANF treatment produced a decrease in fungus viability between 44-42% (p<0.001), only for *C. parapsilosis* C010 isolate the % of fungal damage was less (18 %; p<0.01). For both *C. albicans* clinical isolates used, ANF was ineffective to produce damage to fungal cell forming biofilms regardless of intrinsic ability of this strain (SBF or MBF capacity) (Figure 1).

### Effects of ANF and human monocytes on Candida spp biofilm

In order to explore the ability of immune cells to reduce the fungal biomass of biofilm, we evaluated the capacity of human monocytes alone or in combination with ANF and compared with basal condition and with the effect of anidulafungin treatment alone. For this purpose, mature biofilms of *C. albicans* and *C. parapsilosis sensu stricto* clinical isolates were exposed to three different treatments: a) ANF, b) THP-1 monocytic cell line, c) ANF+THP-1 cells, during 22h. After that, the immune cells were lysed and fungal cell viability was evaluated. Untreated biofilm of each strain was used as growth control. As the resistance of biofilm is associated with the species, the intrinsic capacity of each strain to form biofilm and fungal biomass of biofilm, we performed experiments using biofilm with low (LB) and high biomass (HB) of *C. albicans* and *C. parapsilosis* and exposed them to different treatment (Figure 2 y 3).

**Figure 2.**
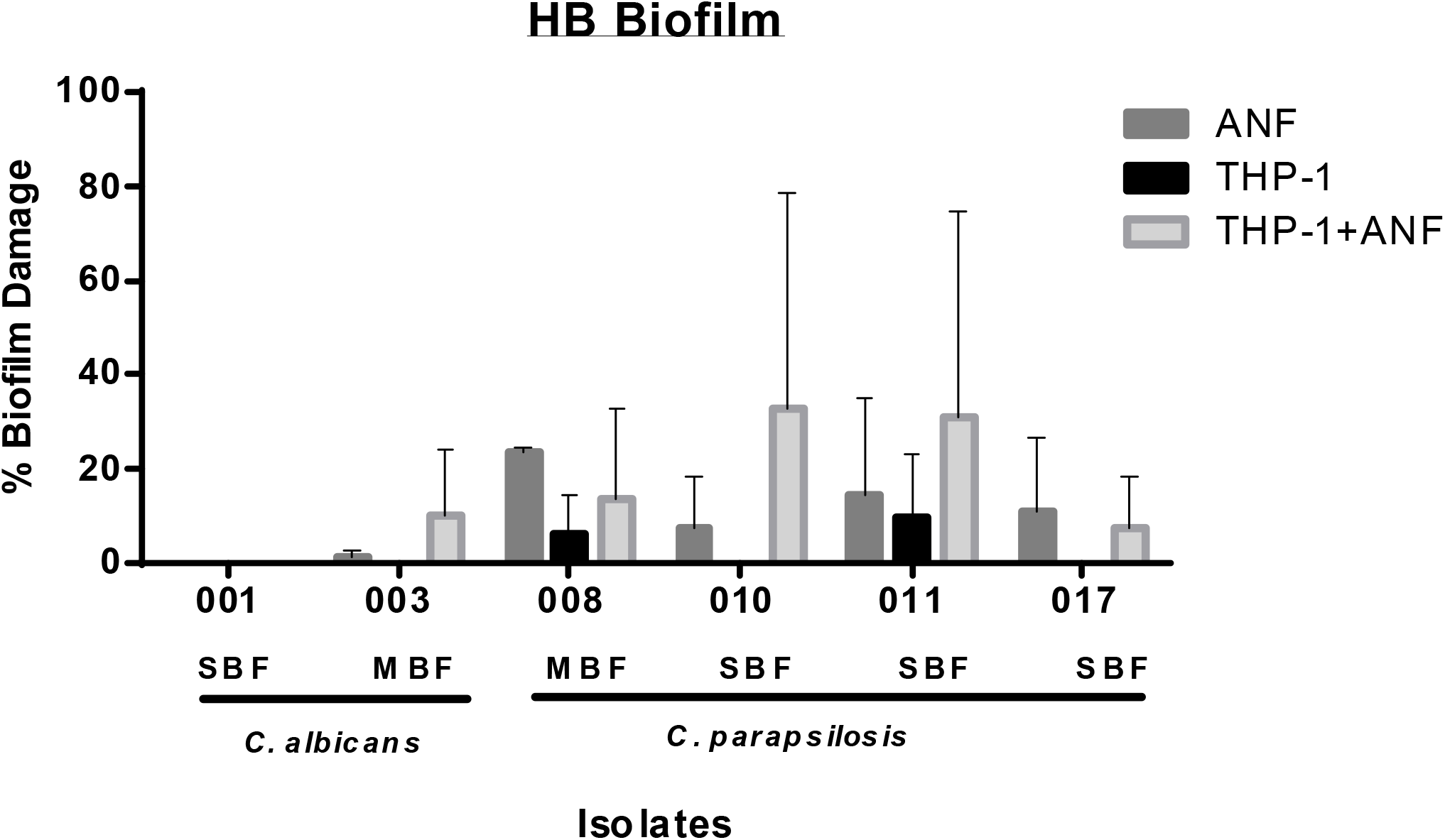
Percentage of damage in high biomass biofilms with different treatments. In order to obtain mature biofilm whit high biomass, 3×10^6^ blastoconidia of each isolate of *C. albicans* or *C. parapsilosis* was cultured for 48 h at 37°C. The percentage of damage induced by ANF (0.12mg/ml), or THP-1 cells (at a 1:1 E:T ratio) or combined treatment against mature *Candida* spp biofilms was evaluated after incubation at 37°C for 22 h. Later THP-1 cells were lised and fungal cell viability measured by XTT assay. The values are means ± SD of three independent experiments. Each experiment was conducted using triplicate wells for each condition. The results of the percentage of damage induced by different treatment was compared with to untreated fungal cell viability (* p<0.01; **p<0.001).

For HB biofilm of the two clinical isolates of *C. albicans* (C001 and C003) all treatments used were unable to modified the fungal biomass, and no changes were observed in fungal viability (Figure 2). In LB mature biofilm, ANF alone or monocytes alone were unable to produce fungal damage, only the combined treatment of immune cells and ANF could be able to damage biofilm compared with untreated control (p<0.01) (Figure 3). Interestingly, a synergistic effect was observed when the treatment with ANF alone was compared with combined treatment for each strain (p<0.01 and p<0.001).

**Figure 3.**
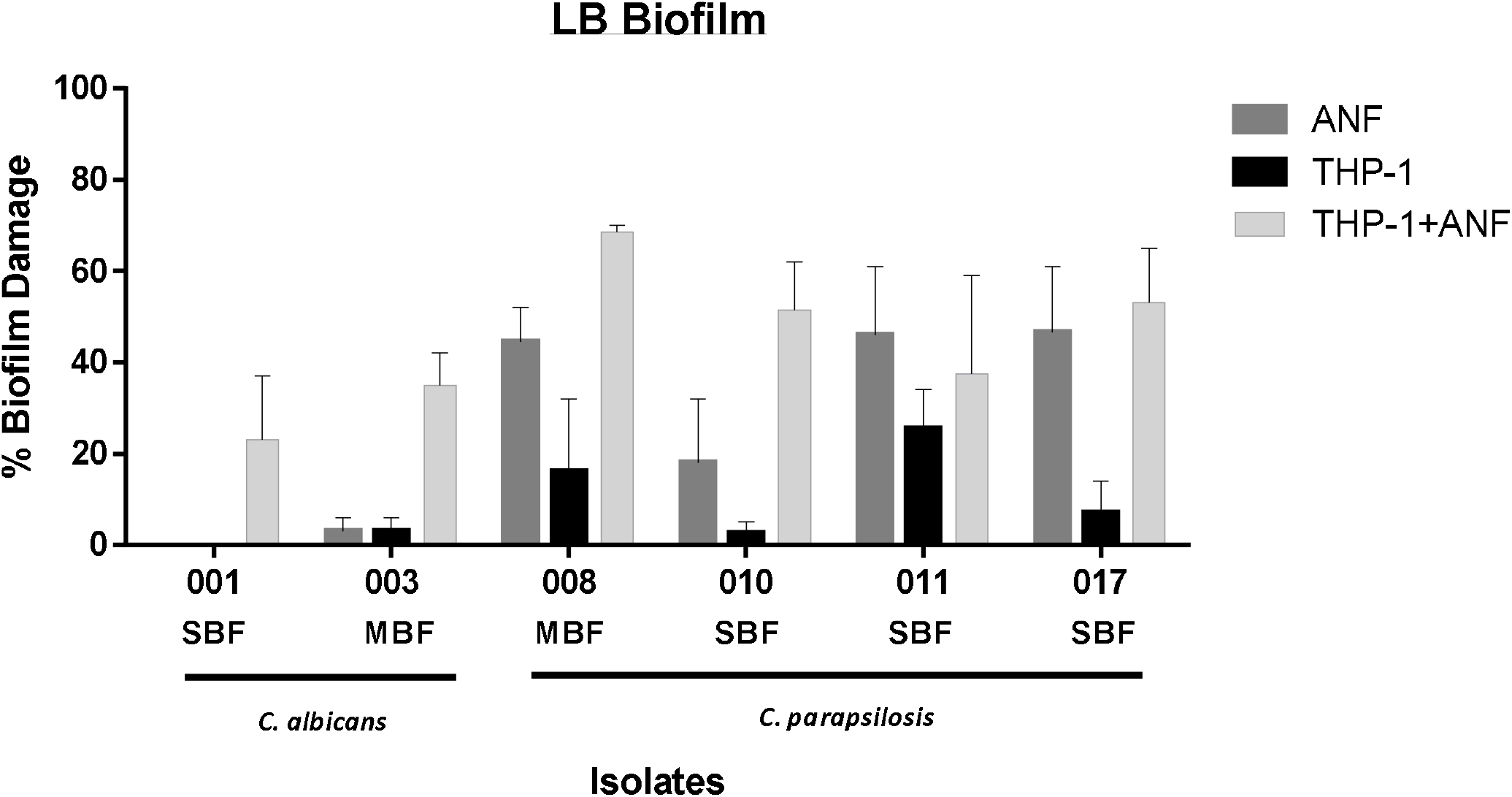
Percentage of damage in low biofilms with different treatments. In order to obtain mature biofilm with low biomass, 1×10^5^ blastoconidia of each isolate of *C. albicans* or *C. parapsilosis* was cultured for 48 h at 37°C. The percentage of damage induced by ANF (0.12mg/ml), or THP-1 cells (at a 1:1 E:T ratio) or combined treatment against mature *Candida* spp biofilms was evaluated after incubation at 37°C for 22 h. Later THP-1 cells were lised and fungal cell viability measured by XTT assay. The values are means±SD of three independent experiments. Each experiment was conducted using triplicate wells for each condition. The results of the percentage of damage induced by different treatment was compared with to untreated fungal cell viability (* p<0.01; **p<0.001). The synergic effect of combined effect of ANF and Mo vs ANF also is indicate (# p<0.01; ## p<0.001).

In *C. parapsilosis sensu stricto* isolates, human Mo were unable to damage all biofilm with HB, and ANF only was effective against C008 strain, which was the only *C. parapsilosis clinical* isolate classified as MBF capacity (p<0.01). The combined treatment showed to be effective over 2/4 biofilms of *C. parapsilosis* isolates (C010 and C011; p<0.01) (Figure 2).

In LB biofilm of *C. parapsilosis sensu stricto* isolates, antifungal treatment with ANF was efficient in 3/4 isolates (C008, C011, C017, p<0.01), meanwhile monocytes alone showed a reduction of biofilms in a range between 4-25%, but this decrease was only significant in C011 isolate (p<0.01). The immune cells in combination with ANF could produce fungal damage in the biofilms of all isolates compared with untreated biofilm (C011 p<0.01; C008, C010 and C017 p< 0.001) (Figure 3) and a synergic effect between the treatments was observed in C008 and C010 isolates (p<0.01). These results demonstrate that while the treatment with ANF alone could be effective to reduce fungal viability of mature biofilms of *C. parapsilosis* with LB (3/4 isolates tested) in a range between 45.5 - 40.0 %, the antifungal agent associated with human Mo, maintained or enhanced this effect in biofilm.

## Discussion

Biofilms are microbial communities embedded in a self-produced extracellular matrix attached to an inert or superficial cellular surface^10,24^. The presence of medical devices such as central venous catheters are known to be important risk factors for the development of biofilm, suggesting that biofilm formation is a key feature in the pathogenesis of candidaemia^25,26^. That is why *Candida* spp. biofilm formation is one of the most extensively investigated virulence factors. Infections related to biofilm are difficult to treat, because antimicrobial resistance is very high in biofilm-producing *Candida* strains ^27–29^. Also infections related to biofilm are difficult to treat, because antimicrobial resistance is very high in biofilm-producing *Candida* strains^26,27^.

*C. albicans* and *C. parapsilosis* exhibited different biofilm architecture and characteristics depending on the ability of each species to produce EPS and display dimorphic growth ^25^. Also, the transcriptional control over processes of biofilm formation like adhesion, filamentation, and EPS production, shows distinctive structure^10^. We included clinical isolates of both species, with different ability to form biofilm, and performed biofilm with two diverse biomasses, high and low, in order to hypothesize they capacity of forming robust biofilms with different behaviors. In this scenario, we evaluate the effects of subinihibitory dose of ANF, human-Mo alone or the combined treatment.

In agreement with previous publications ^21^, biofilms of the clinical isolates evaluated were more resistant than planktonic cells to ANF. About the effect on biofilm, ANF was less effective against HB biofilms than LB of *C. parapsilosis sensu stricto* isolates, and our results also indicated that the antifungal agent used was less effective against *C. albicans*, evidencing a major resistance of this species in concordance with higher pathogenic potential ^30^. These observations may have implications during patients’ treatment, because it has been demonstrated that inadequate antifungal therapy and the presence of an indwelling venous catheter were key predictors of patients mortality and hospital length of stay in patients infected with biofilm forming isolates ^26,31^.

Phagocytes represent the principal effector innate immune in the host to mount an efficient tissue reaction to control the fungus and limit its growth. The reactive oxygen species and nitric oxide produced at high concentrations are directly involved in the killing of this microorganism^8^. Working with THP-1 Mo we observed candidicidal activity against *C. albicans* clinical isolates (data not shown), confirming the ability of this immune population to control the fungus growth. However, the fungus can modulate the immune response favoring its survival and permanence. As MΦ, Mo are a versatile population that vies between two metabolic and functional profile, M1 associated with host protection, and M2 with fungus resistance, making the M1/M2 balance, a relevant event in the development of infection^9^. Working with U937 cell line, we reported that *C. albicans* drives human-Mo toward M2, and decreased IL-1β, IL-8 and TNF levels^9^. In experiments with purified fungal Lip we reported its ability to modulate L-arginine pathway, favored M2 activation in *C. albicans*-primed *in vivo* or *in vitro* MΦ^8,32^. It’s interesting that all clinical isolates evaluated showed Lip activity. Also Chandra *et al*. working with adherent PBMCs and planktonic or *C. albicans* biofilm, showed the relevance of balance between inflammatory/anti-inflammatory mediators in the outcome of *C. albicans*-host immune relationship^33^. We observed that human-Mo alone were unable to induce fungal damage on HB biofilm from *C. albicans* and *C. parapsilosis sensu stricto* clinical isolates, independent of their intrinsic ability to show SBF and MBF capacity. In biofilm with LB of both species, we only observed significant activity in one of *C. parapsilosis* strain.

In addition to well-known antifungal activity, the immunostimulatory properties of many members of echinocandins have been reported. Preexposure of Mo and MΦ to caspofungin(CAS) enhanced fungistatic activity of these cells^34^.

Simitsopoulou *et al*. reported that subinhibitory concentrations of micafungin(MFG) significantly enhanced the expression of innate receptors and IL-1β in macrophage-derived THP-1 exposed to *C. albicans* and *C. parapsilosis* biofilm ^22^. In combined experiments with PMN, macrophage-derived THP-1 and *Candida* biofilm, they showed that the antifungal and immunomodulatory action of pretreatment with MFG has a beneficial effect on immune cells and in control of fungal growth. In the same way, CAS could mediated exposure of fungal β-glucan, the most relevant ligand of Dectin-1^14,22^, suggesting that β-glucan unmasking is probably an echinocandin class effect involved in Mo/MΦ activation. Here, we evaluate the simultaneous effect of subinhibitory doses of ANF and human-Mo on *C. albicans* and *C. parapsilosis* biofilm. The most important contribution of the present study is the evidence that shows the ability of combined treatment of ANF and human-Mo to diminish the mature biofilms with LB in a high proportion of the clinical isolates evaluated.

Although ANF could not appear to achieve complete sterility of *C. albicans* and *C. parapsilosis* in biofilms, it shows synergic effect with immune cells, and can be able to reduce the fungal biomass in established mature biofilm of both species. Interestingly, for *C. albicans* isolates, the independent treatment was not able to reduce biofilm, but the combination, was effective. Also Katragkou *et al*. working with mature biofilm of one isolate of *C. parapsilosis* recovered from sputum, reported that ANF treatment is associated with a significant increase in phagocyte-mediated damage of biofilm^21^. The experimental approach used in our work proposes a more real scenario of study that includes strains with different intrinsic capacity to form biofilm, mature biofilm with LB and HB and structural differences and lays the foundation for upcoming research on this topic. Future treatments that take advantage of the improved Mo/MΦ function, combined with classical or new antifungal drugs constitute a promising strategy to limit formation or to eradicate mature biofilms.

## Acknowledgments

GDV Castillo and AI Azcurra for assistance.

## Author Contributions

Conceptualization, P.A.I. and C.E.S.; methodology, P.A.I, S.C.A, E.R, C.V., M.S.M, C.P and C.A.; formal analysis, P.A.I, S.C.A, E.R and C.E.S.; investigation, F.O.R, J.P.C and C.E.S.; resources, F.O.R, J.P.C and C.E.S; writing-original draft preparation, P.A.I. and C.E.S.; writing-review and editing, P.A.I, S.C.A, E.R, C.V., M.S.M, C.P, C.A., F.O.R, J.P.C and C.E.S; supervision, C.E.S; project administration, C.E.S.; funding acquisition, C.E.S. All authors have read and agreed to the published version of the manuscript.

